# onsite: An Integrated Framework for Phosphosite Localization and False Localization Rate Estimation

**DOI:** 10.64898/2026.07.08.737157

**Authors:** Qi-Xuan Yue, Zhongchun Wei, Chengxin Dai, Mingze Bai, Yasset Perez-Riverol, Timo Sachsenberg

## Abstract

With the rapid development of mass spectrometry-based proteomics, the volume of phosphoproteomic data has increased substantially. However, accurate localization of phosphorylation sites and standardized statistical validation remain critical analytical bottlenecks. To address the lack of standardized cross-algorithm evaluation, we introduce onsite, a unified and open-source Python framework. onsite integrates an alanine-decoy strategy to estimate the false localization rate (FLR) across three algorithms: AScore, PhosphoRS, and pyLucXor. This modular architecture efficiently processes large-scale datasets and enables global FLR calculation. Benchmarking on the standard synthetic phosphopeptide dataset PXD000138 highlighted distinct inter-algorithmic variations. Using the same 5% global FLR threshold, pyLucXor localized the most target sites (28,353). It also reached a high accuracy (91.22%) against the known ground truth, resulting in the largest number of correctly localized sites (25,865). Reanalysis of the highly fractionated, large-scale PXD012255 dataset further demonstrated that native integration of onsite into the quantms pipeline enables scalable processing and provides a standardized framework for FLR control in large-scale phosphoproteomics.

**Graphical Abstract:** 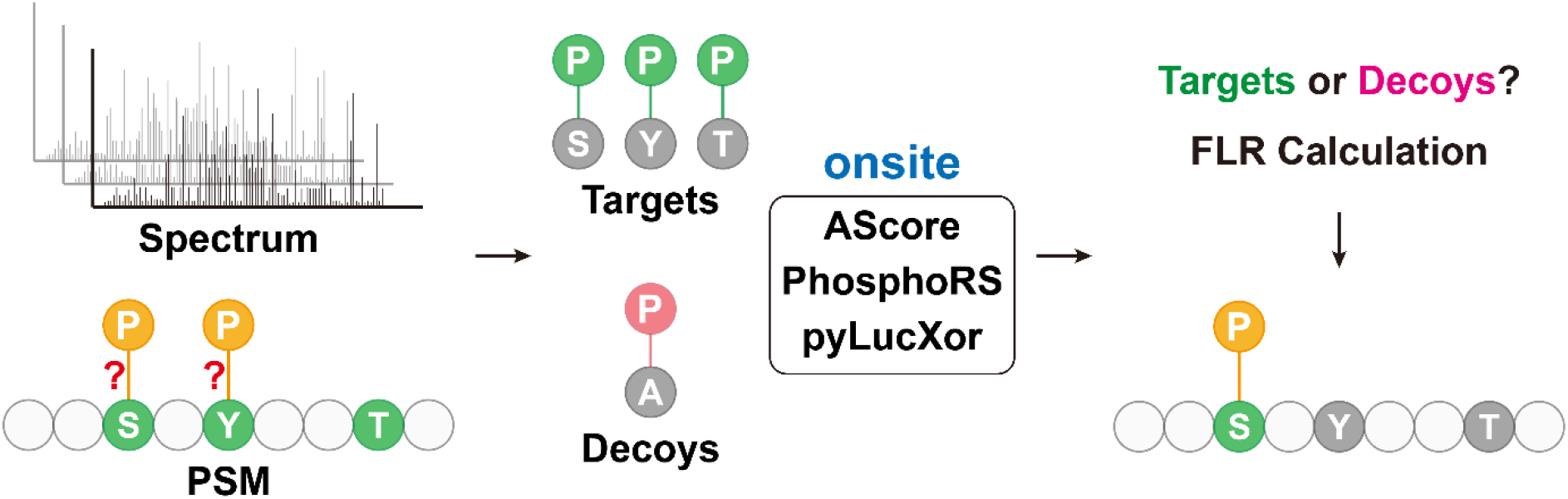

## Introduction

Post-translational modifications (PTMs), particularly phosphorylation, dynamically regulate protein structure, function, and cellular signaling^1, 2^. Phosphorylation occurs primarily on serine (S), threonine (T), and tyrosine (Y) residues^3^. However, peptides often contain multiple candidate phosphorylation sites, making precise localization of the modification to a specific residue a critical step in phosphoproteomic analysis. Accurate identification of these sites is essential for deciphering the molecular mechanisms governing protein function and regulatory networks.

While standard peptide database searches can match tandem mass spectrometry (MS/MS) spectra to peptide sequences, they are not primarily designed to accurately localize post-translational modifications. They provide confidence estimates for peptide-spectrum matches (PSMs) but typically lack a dedicated assessment of modified site localizations^4^. This poses a particular challenge for peptides containing multiple candidate residues, where the correct phosphate-group localization remains ambiguous. To resolve positional uncertainties, the need for accurate site localization has driven the development of specialized algorithms. Early pioneering methods utilized probability-based models. For instance, the PTM score^2^ calculates site-specific localization probabilities to classify high-confidence phosphosites and was subsequently integrated into MaxQuant. Similarly, the AScore algorithm employs a binomial statistics metric based on site-determining ions to automate localization^5^. As a computationally less demanding approach, the Mascot Delta Score infers localization confidence from the difference in search-engine scores between the most likely and the next-best phosphorylation-site assignment^6^. PhosphoRS uses probabilistic scoring to analyze isoforms through a peak extraction technique adaptable to various fragmentation methods^7^. Because neither probability-nor delta-scoring methods provide direct control of localization error rates at the dataset level, LuciPHOr incorporated a modified target–decoy strategy. Using decoy peptides with artificial modifications on non-candidate residues, it enables simultaneous localization and directly estimates the false localization rate (FLR)^4^. Its successor, LuciPHOr2, extends this FLR estimation to generic PTMs with significantly reduced computation time^8^.

To establish a universally comparable metric for FLR estimation, the phospho-Alanine (pAla) decoy strategy was developed^9^. Unlike methods that depend on algorithm-specific localization scores, pAla introduces decoy site assignments by allowing phosphorylation on alanine residues, enabling direct estimation of localization error rates. Alanine provides a chemically implausible decoy site for conventional phosphoproteomics. However, target (S/T/Y) and decoy (A) residues naturally occur at different background frequencies within the proteome. Consequently, accurate FLR estimation requires statistical calibration to account for differences in residue frequencies. After calibration, the pAla decoy method enables consistent and method-independent estimation of the FLR across datasets and localization algorithms.

We present onsite, an open-source implementation of alanine-decoy-based FLR estimation that is fully compatible with OpenMS^10^ and quantms^11^. Rather than introducing a new localization model, onsite provides a unified framework for applying the pAla decoy strategy across established localization algorithms, including AScore, pyLucXor (a Python reimplementation of LuciPHOr2), and PhosphoRS. While the core module performs FLR estimation on individual files, dedicated utility scripts enable aggregation of statistics across multiple files facilitating large-scale phosphoproteomics analysis. By integrating pAla-based FLR estimation into modern bioinformatics workflows, onsite provides a common framework for translating algorithm-specific localization scores into universally comparable estimates of dataset-wide localization error.

### Experimental Procedures

#### Implementation

Built on PyOpenMS, onsite implements a modular pipeline that spans data import, peptide-spectrum matching, site localization, and FLR calculation, accepting input in open formats. To support large-scale analyses, it parallelizes computationally intensive tasks and aggregates target and decoy observations across files for dataset-level FLR estimation. onsite implements the pAla target-decoy approach (TDA), in which alanine serves as a decoy to model the distribution of random mislocalizations. onsite integrates three localization algorithms. The first algorithm, AScore, localizes phosphorylation sites by comparing alternative site assignments and assessing the evidence provided by site-determining fragment ions. Using a binomial probability model, it calculates how strongly the spectrum supports one localization over competing possibilities under different levels of peak filtering. Higher scores indicate a lower probability of incorrect site assignments. The second algorithm, PhosphoRS, evaluates all feasible site localizations within a probabilistic framework and outputs posterior probabilities for individual modification sites. This allows PhosphoRS to provide a more direct and interpretable measure of site-specific localization confidence compared to AScore. Third, pyLucXor introduces a rigorous delta score distribution model.

Finally, a global pAla FLR module calculates and aggregates statistics across multiple files. It excludes unambiguous peptides (where all potential sites are modified) to prevent an artificial underestimation of the global FLR. onsite reports localization results using the standardized ProForma notation^12^, ensuring interoperability.

#### Benchmarking

We evaluated onsite using the synthetic phosphopeptide library dataset PXD000138, comprising 96 HCD raw files available through PRIDE. Peptide identifications generated with the quantms workflow were used as input for a standardized comparison of phosphorylation-site localization algorithms^13^. The three algorithms (AScore, pyLucXor, and PhosphoRS) implemented in the onsite package were used for the comparative analysis (**Table S-1**).

Initially, we systematically compared the localization sites predicted by the algorithms under unified, standardized threshold conditions. To ensure an unbiased evaluation and to strictly isolate localization accuracy from sequence identification confounding factors, we implemented a rigorous, uniform filtering pipeline across all algorithms, integrating principles from Ramsbottom et al. ^9^ and Camacho et al. ^14^:

The unified filtering pipeline comprised four sequential steps. First, all decoy peptide-spectrum matches were removed and FDR was controlled by applying a strict PSM-level q-value threshold of < 0.01. Second, to eliminate background noise, non-synthetic peptides were excluded, and only peptides that perfectly matched the target sequences within the synthetic seed library were retained. Third, to eliminate the confounding complexity of co-occurring modifications, multi-site-phosphorylated peptides were excluded. Finally, unambiguous peptides (i.e., those with no alternative localization options) were discarded, with the strict prerequisite that a reported site be included in the analysis only if it has a usable localization confidence score.

We estimate the FLR according to Camacho et al. ^14^ on the sorted site scores:

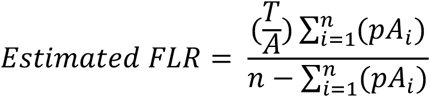

where: *T* and *A* represent the counts of target and decoy amino acids, 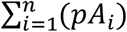 represents the cumulative number of decoy sites, and n represents the rank.

Secondly, we evaluated localization accuracy against a ground-truth reference derived from the synthetic phosphopeptide library. Known phosphorylation sites reported in Supplementary Table 2 of the original publication served as the reference set^13^. After applying the same quality-control criteria described above and filtering results at an estimated global FLR of 5%, site assignments were compared with the ground truth to determine true-positives and false-positives for each method. We calculate the true FLR, using the method proposed by Camacho et al. ^14^:

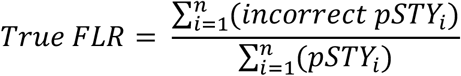

Additionally, we evaluated a conventional database search engine as a baseline using the same filtering criteria as the onsite benchmark. Restricting the analysis to single-site phosphopeptides eliminated localization ambiguity, allowing peptide identification scores to be directly interpreted as localization confidence.

#### Interoperability with quantms and support of HUPO-PSI standards

By default, quantms performs site localization using pyLucXor with its standard parameter settings (default parameters shown in **Table S-3** and **Table S-4**). Users can alternatively select AScore or PhosphoRS through the configuration file (both defaulting to a fragment mass tolerance of 0.05 Da) and customize algorithm-specific parameters as needed. Supported input formats include the community-standard HUPO-PSI formats mzML for spectral data (-in) and mzIdentML^15^ for peptide-spectrum matches (-id), together with the OpenMS-native idXML format and the columnar idparquet format (QPX, https://github.com/bigbio/qpx).

## Results and Discussion

pyLucXor could not process three low-yield files (73, 94, and 96) because they contained too few high-confidence PSMs for model training. To maintain a consistent comparison across methods, these files were excluded from all analyses, despite being processable by AScore and PhosphoRS. Of the 336,993 initial PSMs, 265,648 were common to all three localization tools. After filtering, including the exclusion of unambiguous peptides, approximately 84,100 PSMs were retained for downstream analysis. **Figure 1A** shows localization performance across a range of global FLR thresholds. AScore and PhosphoRS identify more target phosphorylation sites at low FLRs (e.g., 1%), whereas pyLucXor achieves the highest yield as the FLR threshold becomes less stringent (e.g., 5% and 10%).

**Figure 1.**
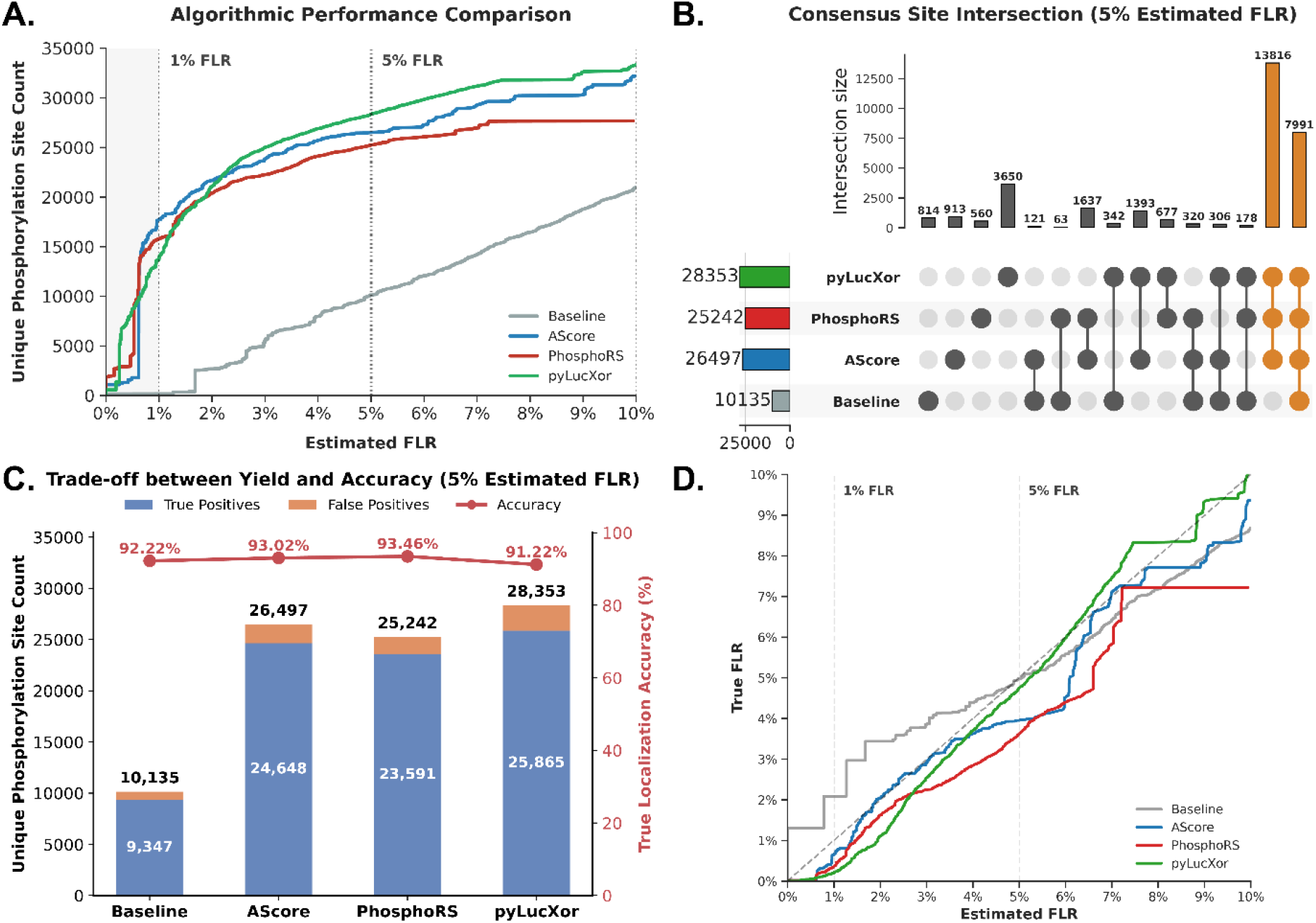
Performance comparison of phosphorylation site localization algorithms under standardized global FLR control. (A) Number of localized target phosphorylation sites reported by the baseline approach, AScore, PhosphoRS, and pyLucXor as a function of estimated global FLR. Dashed lines indicate the 1% and 5% FLR thresholds commonly used for downstream analyses. (B) UpSet plot showing the overlap of target site localizations at a 5% estimated global FLR. The majority of sites were shared among algorithms, while pyLucXor contributed the largest set of unique localizations. Numbers to the left indicate the total number of target sites localized by each method. (C) Comparison of localization yield and accuracy at 5% estimated global FLR. Bars show the numbers of true-positive and false-positive site localizations determined using the synthetic peptide ground-truth dataset, while the red line indicates localization accuracy. pyLucXor achieved the highest site yield, whereas PhosphoRS achieved the highest localization accuracy. (D) Calibration of estimated versus observed (true) FLR. Perfect calibration corresponds to agreement between estimated and true FLR.

At a global FLR threshold of 5%, pyLucXor identified the largest number of target sites (**Table 1**), recovering 28,353 localizations compared with 26,497 for AScore and 25,242 for PhosphoRS. To further characterize these results, we examined the overlap and method-specific localizations at the 5% FLR cutoff **(Figure 1B**). UpSet analysis showed that 13,816 sites were identified by all three algorithms, whereas pyLucXor uniquely localized 3,650 sites not reported by either AScore or PhosphoRS. These results indicate that pyLucXor’s increased yield is largely driven by additional site recoveries beyond the consensus set, while decoy rates remained comparable across methods (6.57–12.57%).

**Table 1.** Localization performance of the evaluated algorithms at fixed global FLR thresholds. Numbers of localized target phosphorylation sites identified at 1%, 5%, and 10% global FLR. Localized target and decoy-A sites across the full dataset, with the decoy rate in parentheses.

| Algorithm | 1% FLR | 5% FLR | 10% FLR | Total sites/decoys (rate) |
| --- | --- | --- | --- | --- |
| **pyLucXor** | 13,892 | 28,353 | 33,372 | 39,762 / 3,374 (8.49%) |
| **AScore** | 17,727 | 26,497 | 32,195 | 38,198 / 2,508 (6.57%) |
| **PhosphoRS** | 15,819 | 25,242 | 27,682 | 40,068 / 5,035 (12.57%) |
| **Baseline** | 180 | 10,135 | 21,015 | 40,155 / 3,763 (9.37%) |

To assess the balance between localization yield and accuracy, we compared the numbers of true-positive (TP) and false-positive (FP) phosphorylation site localizations at a global FLR threshold of 5% (**Figure 1C**). The baseline search-engine approach identified 9,347 TP sites, highlighting the limited ability of peptide-level confidence scores to resolve ambiguous modification sites. In contrast, the dedicated localization algorithms substantially increased the number of correctly localized sites, with AScore, PhosphoRS, and pyLucXor identifying 24,648, 23,591, and 25,865 TP sites, respectively. PhosphoRS and AScore achieved the highest localization accuracies (93.46% and 93.02%, respectively), whereas pyLucXor recovered the most correctly localized sites at a marginally lower accuracy, keeping it within about two percentage points of the best-performing method (91.22% versus 93.46%), reflecting a favorable balance between localization sensitivity and accuracy.

We evaluated algorithm calibration by comparing estimated and observed FLRs (**Figure 1D**). The baseline method underestimated localization error in the low FLR region, whereas PhosphoRS produced conservative FLR estimates throughout the evaluated range. AScore was conservative at low FLRs but became less conservative at higher thresholds. pyLucXor exhibited the closest overall agreement between estimated and observed FLR, although it was slightly conservative below 5% FLR and slightly anti-conservative between 5% and 10% FLR before converging near the 10% threshold. Overall, all localization algorithms yielded reasonably well-calibrated results when using the pAla TDA for FLR estimation.

Computational performance was assessed by processing each dataset file individually and measuring execution time (**Table S-2**). AScore achieved the highest throughput (227.17 PSMs/s), while pyLucXor was the slowest integrated method (66.36 PSMs/s), consistent with its more extensive statistical modeling (**Table S-2**). Nevertheless, pyLucXor outperformed the original Java implementation of LuciPHOr2 (62.1 PSMs/s).

To facilitate adoption and reproducibility, onsite is openly available on GitHub (https://github.com/bigbio/onsite) and distributed via PyPI as a standard Python package. The software provides both an application programming interface (API) and a command-line interface (CLI) for local phosphorylation-site localization analyses.

Furthermore, to ensure platform-independent deployment and reproducible analyses, onsite is distributed through the BioContainers registry. This containerized implementation allows onsite to function as a self-contained module within quantms, a Nextflow-based workflow for quantitative proteomics, thereby integrating multi-algorithm phosphorylation-site localization into automated data-processing pipelines. Through Nextflow’s orchestration capabilities, the module can be deployed reproducibly across local, high-performance computing, and cloud environments, supporting scalable reanalysis of large-scale phosphoproteomics datasets.

To demonstrate the scalability and multi-algorithmic capabilities of our framework, we reanalyze a large-scale phosphoproteomics dataset (PXD012255) of the HCT116 colon carcinoma cell line^16^ using the quantms workflow (v1.7.0). By enabling the onsite_compute_all_scores parameter, AScore, PhosphoRS, and pyLucXor were executed within a single automated workflow. The full reanalysis was performed on a local workstation (Intel Core i9-7900X, 128 GB RAM, Ubuntu 24.04.2 LTS) and completed in 15h 29m 18s.

Following data processing, we assessed algorithm performance across the highly fractionated dataset. Despite the data sparsity and variability introduced by capillary zone electrophoresis (CZE) separation, quantms-onsite successfully generated localization scores for all three algorithms across the cohort (**Figure 2A**). To evaluate the complementarity of the methods, we analyzed the overlap of phosphorylation sites localized at a stringent 1% global FLR after aggregating results across all fractions (**Figure 2B**). UpSet analysis revealed a substantial consensus set of 4,421 sites shared by all three algorithms, alongside a large number of method-specific localizations, including 3,742 sites uniquely identified by pyLucXor. These results demonstrate that the algorithms provide complementary localization evidence and that combining multiple approaches could increase phosphosite coverage beyond that achieved by any individual method.

**Figure 2.**
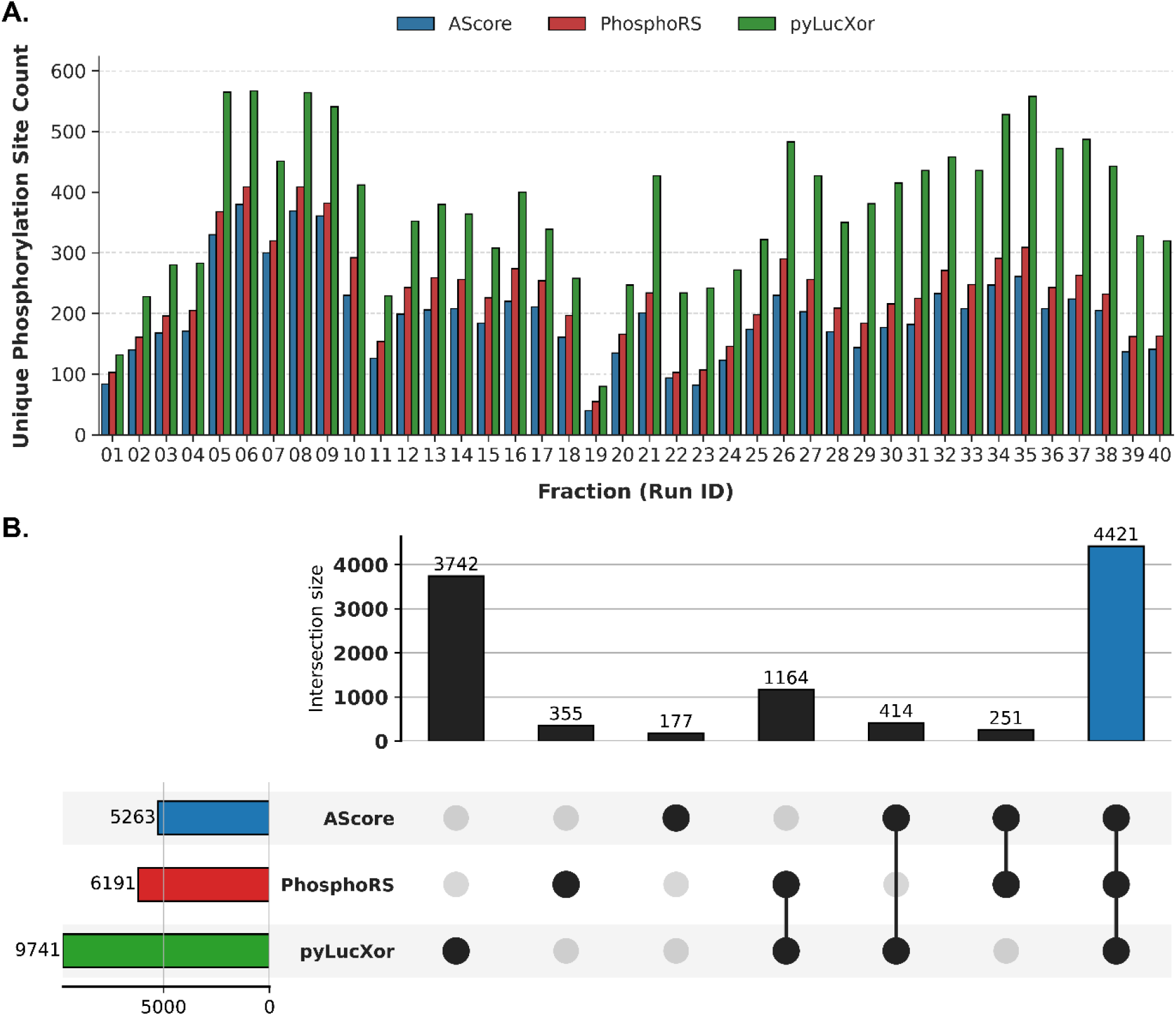
Application of the quantms (onsite) workflow to the HCT116 cell-line dataset (PXD012255). (A) Per-file distribution of high-confidence phosphorylation sites across 40 runs, filtered at a 1% global FLR threshold. (B) UpSet plot illustrating the overlap and algorithm-specific high-confidence phosphorylation sites identified by AScore, PhosphoRS, and pyLucXor.

## Conclusion

We present onsite, an open-source Python framework that integrates AScore, PhosphoRS, and pyLucXor within a unified phosphorylation-site localization workflow. By harmonizing algorithm inputs and outputs and implementing a standardized alanine-decoy strategy, onsite supports empirical estimation and consistent control of the global false localization rate across multiple LC–MS/MS files.

Independent benchmarking against the known phosphorylation sites in the synthetic PXD000138 dataset confirmed the localization performance of the integrated algorithms while revealing distinct trade-offs between yield and accuracy. At a 5% global FLR, pyLucXor recovered the largest number of correctly localized sites, whereas AScore and PhosphoRS achieved slightly higher localization accuracy. Analysis of the large-scale PXD012255 dataset further revealed substantial agreement among the algorithms together with sizeable method-specific site sets, demonstrating their complementary contributions to phosphosite coverage.

Implemented using PyOpenMS and distributed via PyPI and BioContainers, onsite can be deployed as a standalone tool or as part of the quantms workflow. Its containerized implementation supports reproducible execution across local workstations, high-performance computing systems, and cloud environments. Together, standardized global FLR estimation, independent ground-truth validation, and scalable workflow integration make onsite a practical framework for reproducible, cross-algorithm comparable phosphorylation-site localization in large-scale phosphoproteomics. Future work will focus on improving computational efficiency and extending the framework to additional post-translational modifications.

## ASSOCIATED CONTENT

### Supporting Information

The Supporting Information is available free of charge at https://pubs.acs.org/doi/[DOI]: algorithm comparison and computational performance (Tables S-1 and S-2) and default localization parameters (Tables S-3 and S-4).

### Data Availability Statement

onsite is an open-source Python library (https://github.com/bigbio/onsite). All datasets reanalysed in this study are publicly available in the PRIDE repository under accession numbers PXD000138 and PXD012255.

## AUTHOR INFORMATION

### Author Contributions

Qi-Xuan Yue: Software, Methodology, Validation, Investigation, Visualization, Writing – original draft, Writing – review & editing. Zhongchun Wei: Software. Chengxin Dai: Software, Data curation, Methodology, Writing – original draft. Mingze Bai: Validation, Resources, Supervision, Funding acquisition, Writing – review & editing. Yasset Perez-Riverol: Conceptualization, Software, Methodology, Resources, Supervision, Funding acquisition, Writing – original draft, Writing – review & editing. Timo Sachsenberg: Conceptualization, Software, Methodology, Supervision, Project administration, Funding acquisition, Writing – original draft, Writing – review & editing.

## Acknowledgment

ChatGPT and Claude were used for language editing to improve the clarity, readability, and flow of the manuscript. The authors reviewed and edited the AI-assisted text and take full responsibility for the final content.

