## Supplemental Information for "onsite: An Integrated Framework for Phosphosite Localization and False Localization Rate Estimation"

Qi-Xuan Yue [a], Zhongchun Wei [a], Chengxin Dai [b,c], Mingze Bai [a], Yasset Perez-Riverol [d], Timo Sachsenberg [e,f*]
a. Chongqing Key Laboratory of Big Data for Bio Intelligence, Chongqing University of Posts and Telecommunications, Chongqing, China
b. State Key Laboratory of Medical Proteomics, Beijing Proteome Research Center, National Center for Protein Sciences (Beijing), Beijing Institute of Lifeomics, 102206, Beijing, China
c. International Academy of Phronesis Medicine (Guangdong), 510320, Guangdong, China
d. European Molecular Biology Laboratory, European Bioinformatics Institute, Wellcome Genome Campus, Cambridge, United Kingdom
e. Applied Bioinformatics, Department of Computer Science, University of Tübingen, Tübingen, Germany
f. Institute for Bioinformatics and Medical Informatics, University of Tübingen, Tübingen, Germany

**Supplementary Table S-1:** Commands and parameters of the evaluated algorithms.

| Algorithm | Commands and parameters |
|---|---|
| *AScore (onsite)* | onsite ascore -in mzml_path -id idxml_path -out output_path -threads 1 --fragment-mass-tolerance 0.05 --add-decoys |
| *PhosphoRS (onsite)* | onsite phosphors -in mzml_path -id idxml_path -out output_path -threads 1 --fragment-mass-tolerance 0.05 --add-decoys |
| *pyLucXor (onsite)* | onsite lucxor -in mzml_path -id idxml_path -out output_path -threads 1 --fragment-method HCD --fragment-mass-tolerance 0.05 --target-modifications "Phospho(S),Phospho(T),Phospho(Y),PhosphoDecoy(A)" |
| *Luciphor (original)* | LuciphorAdapter -in mzml_path -id idxml_path -out output_path -executable /path/to/luciphor2.jar -target_modifications 'Phospho (S)' 'Phospho (T)' 'Phospho (Y)' -fragment_mass_tolerance 0.05 -fragment_error_units Da -fragment_method HCD -fragment_error_value 0.05 -fragment_error_unit Da -threads 1 -num_threads 1 |

Luciphor: quay.io/biocontainers/openms-thirdparty:3.5.0--h9ee0642_0

**Supplementary Table S-2:** Computational time benchmark and throughput for the evaluated algorithms integrated within the onsite toolkit.

| **Algorithm** | **Execution Time (seconds)** | **Throughput (PSMs/second)** |
|---|---|---|
| *Luciphor (original)* | 5426.65 | 62.10 |
| *pyLucXor (onsite)* | 5078.61 | 66.36 |
| *AScore (onsite)* | 1483.43 | 227.17 |
| *PhosphoRS (onsite)* | 1504.07 | 224.05 |

Note: The benchmark was performed on a workstation with an Intel Core i9-7900X CPU, 128 GB RAM, and Ubuntu 24.04.2 LTS.

**Supplementary Table S-3:** Global pipeline parameters (these parameters are common to all three algorithms).

| Parameter | Description | Default value |
|---|---|---|
| *-in* | Input spectrum file | mzML |
| *-id* | Input identification file | MzIdentML, idXML, idparquet (QPX) |
| *-out* | Output file | - |
| *--fragment-mass-tolerance* | Fragment mass tolerance for the global search | 0.05 |
| *--fragment-mass-unit* | Unit for fragment mass tolerance | Da |
| *--threads* | Number of parallel threads for execution | 1 |
| *--add-decoys* | Add decoy modifications | False |
| *--compute-all-scores* | Run all three localization algorithms and merge the results | - |

**Supplementary Table S-4:** pyLucXor specific parameters.

| Parameter | Description | Default value |
|---|---|---|
| *--fragment-method* | Fragmentation method utilized (CID or HCD) | CID |
| *--fragment-mass-tolerance* | Tolerance for matching fragment ions | 0.5 |
| *--min-mz* | Minimum m/z threshold; peaks below this value are ignored | 150.0 |
| *--target-modifications* | Target modification residues | 'Phospho(S),Phospho(T),Phospho(Y)' |
| *--neutral-losses* | List of target neutral losses | sty -H3PO4 -97.97690 |
| *--decoy-mass* | Mass added for decoy modification generation | 79.966331 |
| *--decoy-neutral-losses* | List of decoy neutral losses | X -H3PO4 -97.97690 |
| *--max-charge-state* | Maximum precursor charge state to consider | 5 |
| *--max-peptide-length* | Maximum allowed peptide length | 40 |
| *--max-num-perm* | Maximum number of permutations | 16384 |
| *--modeling-score-threshold* | Threshold applied for score modeling | 0.95 |
| *--scoring-threshold* | Minimum score threshold | 0.0 |
| *--min-num-psms-model* | Minimum number of PSMs for modeling | 50 |
| *--seed* | Random Number Generator (RNG) seed for reproducible decoy permutations and model subsampling | 42 |
| *--rt-tolerance* | Retention time tolerance | 0.01 |
| *--disable-split-by-charge* | Disable the splitting of scoring models by charge state | False |
